# WebAtlas pipeline for integrated single cell and spatial transcriptomic data

**DOI:** 10.1101/2023.05.19.541329

**Authors:** Tong Li, David Horsfall, Daniela Basurto-Lozada, Kenny Roberts, Martin Prete, John E G Lawrence, Peng He, Elisabeth Tuck, Josh Moore, Shila Ghazanfar, Sarah Teichmann, Muzlifah Haniffa, Omer Ali Bayraktar

## Abstract

Single cell and spatial transcriptomics illuminate complementary features of tissues. However, online dissemination and exploration of integrated datasets is challenging due to the heterogeneity and scale of data. We introduce the WebAtlas pipeline for user-friendly sharing and interactive navigation of integrated datasets. WebAtlas unifies commonly used atlassing technologies into the cloud-optimised Zarr format and builds on Vitessce to enable remote data navigation. We showcase WebAtlas on the developing human lower limb to cross-query cell types and genes across single cell, sequencing- and imaging-based spatial transcriptomic data.

---

Single cell and spatial transcriptomics provide complementary tools for tissue atlassing. While single cell RNA-sequencing (scRNA-seq) profiles whole-transcriptomes of individual cells, it does not capture their spatial locations in tissues. Conversely, sequencing-based spatial transcriptomics (ST) can profile whole transcriptomes *in situ*, but widely used technologies such Visium and Slide-Seq do not provide single cell spatial resolution. Although alternative imaging-based ST methods such as In Situ Sequencing and MER-FISH provide true single cell resolution, they are limited to targeted gene panels of around 100 to 1000 genes^1^.

Computational integration can harness the strengths of scRNA-seq and ST modalities to resolve cell types and impute transcriptomes of single cells *in situ*^2^. However, we lack user-friendly software solutions to disseminate and navigate integrated single cell and spatial transcriptomic datasets. First, scRNA-seq and ST data objects are often saved in non-unified sequencing and imaging file formats that perform poorly with web technologies^3–5^. Second, existing software platforms do not readily support simultaneous browsing of multiple integrated data modalities^3, 5, 6^. These limitations impede usable and interpretable access to a wealth of tissue atlas data for the research community, and hinder biological insights that can only be gained from integrated datasets.

Here, we introduce the WebAtlas pipeline to enable online sharing and navigation of integrated single cell and spatial transcriptomic datasets (Fig 1A). We address the aforementioned challenges with two main innovations. First, we provide a new data ingestion pipeline to convert and unify datasets from multiple single cell and spatial technologies into the cloud-ready Zarr format^7^. Second, we provide a frontend web client based on the Vitessce framework^8^ to enable interactive exploration of single cell and spatial data through a web browser, where users can intuitively cross-query gene expression and cell types across modalities (Fig 1A). WebAtlas enhances usability, interpretability and equitable access of multi-modal tissue atlases, facilitating biological insight for a global audience (see Sup Note 1 for detailed comparison to other platforms).

**Figure 1.**
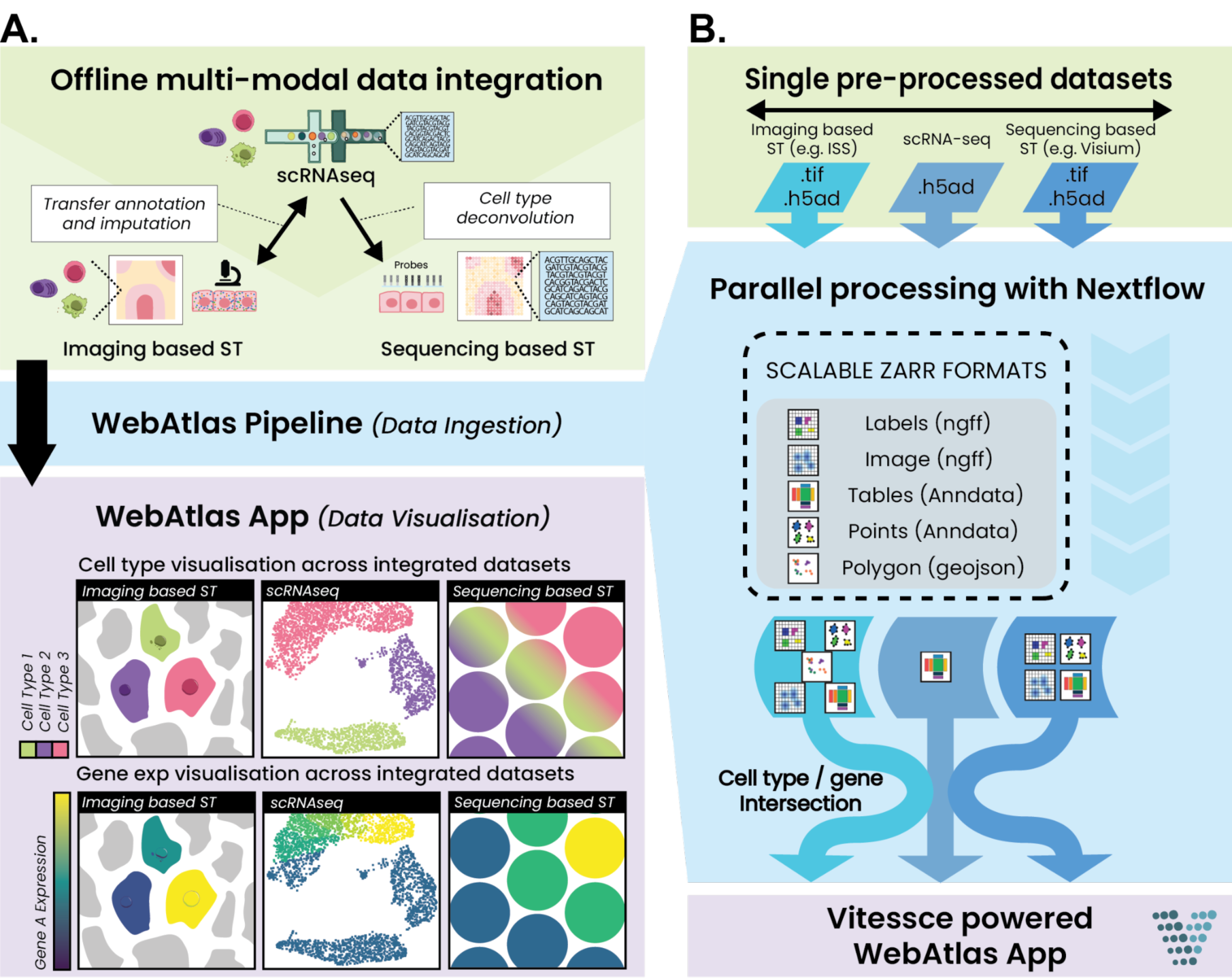
Overview of WebAtlas pipeline. A. WebAtlas incorporates integrated scRNA-seq, imaging- and sequencing-based ST datasets for interactive web visualisation, enabling cross-query of cell types and gene expressions across modalities. B. WebAtlas data ingestion pipeline converts diverse data objects from integrated single cell and spatial technologies to the Zarr format and runs on Nextflow. The pipeline extracts shared genes and cell types features across modalities.

On WebAtlas, single cell and spatial datasets are linked by biomolecular metadata, such as shared cell type or gene annotations. Linkage is performed prior to WebAtlas ingestion using existing data integration methods like Cell2location^9^ and StabMap^10^ that map scRNA-seq cell type references onto ST datasets and impute unobserved gene expression in the latter (Fig 1A).

Our re-usable data ingestion pipeline performs the extract-transform-load steps for sequencing and imaging data objects generated from various single cell and spatial transcriptomic technologies (Fig 1B). To enable scalable and interoperable online browsing of tissue atlas datasets, our pipeline produces a standardised output using the Zarr file format, which utilises an array chunking strategy for efficient data access from the cloud. Our Zarr convention is closely aligned with the recently released SpatialData format^11^, supporting data standardisation efforts.

Tabular gene expression and cell type annotation files for scRNA-seq and ST are converted from the commonly used AnnData format^12, 13^ into AnnData-Zarr. The raw image components of ST data, including fluorescent or brightfield microscopy images along with label images from cell segmentation, are converted into OME-Zarr format^14^ to enable efficient multi-scale visualisation. Other ST data elements, such as points (*e.g.* RNA spots/molecules) and polygons (*e.g.* cell masks), are also stored in Zarr according to the AnnData specification. Together with the authors of SpatiaData, we propose that these structures are included in the community-defined next-generation file format (NGFF)^15^ to achieve a FAIR^16^ exchange of multi-modal tissue atlas data.

The WebAtlas data ingestion pipeline is implemented on Nextflow and is configured through a simple YAML schema that defines input data files and visualisation parameters. Flexibility to support a diverse set of use-cases is built into the pipeline. Input datasets are processed independently and users can choose to process an individual dataset (*e.g.* Visium), specific components of a given dataset (*e.g.* cell segmentation masks but not raw images) or any given combination of integrated datasets (*e.g.* scRNA-seq and Visium or scRNA-seq and imaging). This modular design also supports extensions to new data modalities and formats, such as customised microscopy (see below). The pipeline outputs Zarr formatted datasets and configuration files for web visualisation.

To visualise and cross-query integrated datasets, we use the open-source Vitessce tool. Vitessce provides a customisable and serverless web framework for interactive exploration of single cell and spatial data. However, it has not been utilised to date on fully integrated single cell and spatial modalities, and its configuration requires programmatic expertise. To coordinate Vitessce on computationally integrated datasets, the WebAtlas pipeline first identifies shared ontology features such as genes and cell types across all modalities. To simplify visualisation through Vitessce, the pipeline generates a default View Config JSON file, which instructs Vitessce to access pertinent data features from each ingested modality (Sup Methods). Each Vitessce component is bound to a specific dataset and all components are updated in a coordinated manner to visualise queried genes or cell types (Fig 2A).

**Figure 2.**
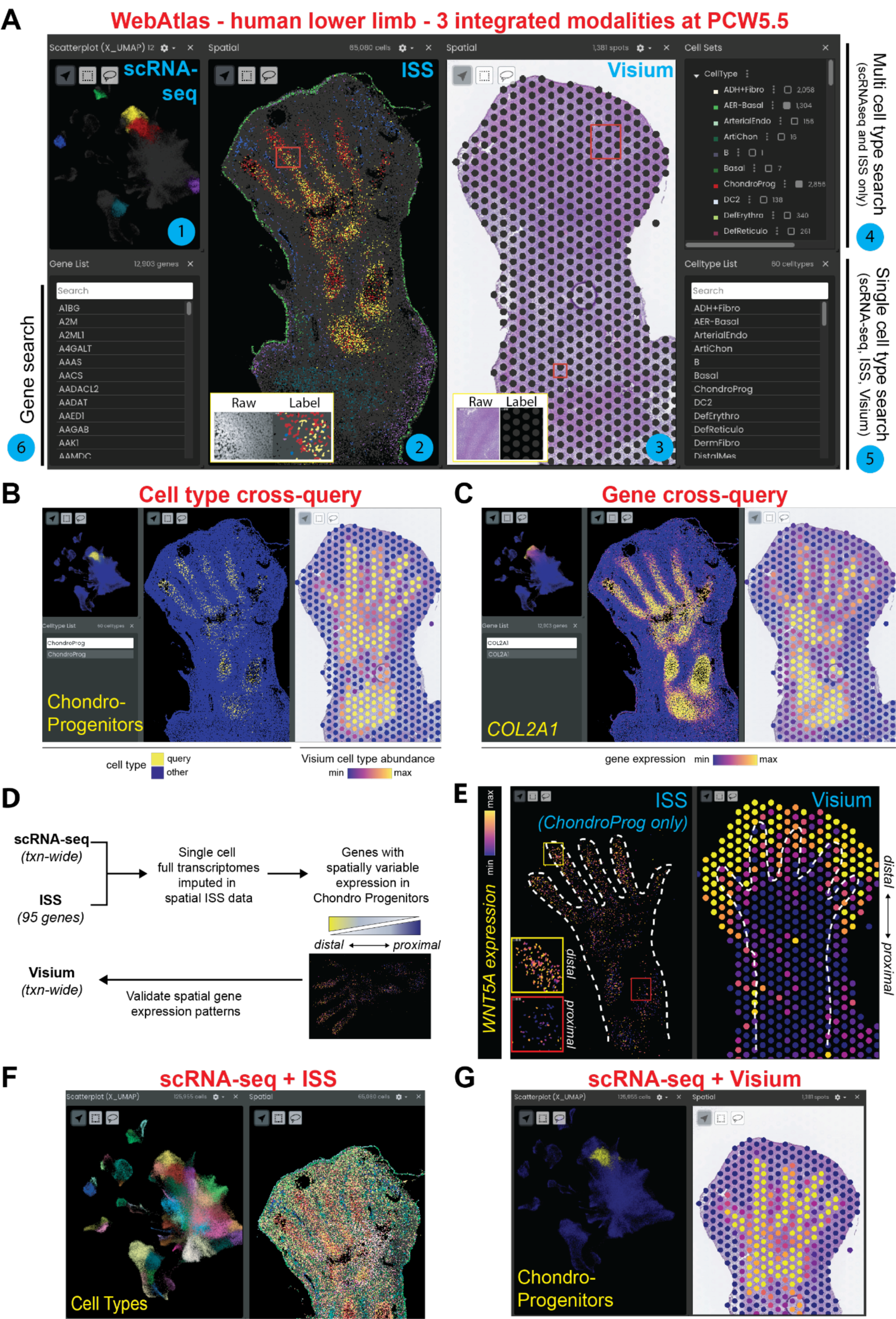
The human lower limb WebAtlas. A. A snapshot of the WebAtlas app visualising integrated scRNA-seq, ISS and Visium datasets of the human lower limb at 5.5 post conception weeks (PCW5.5). The Vitessce component windows show the following as numbered. (1) UMAP representation of cell types in scRNA-seq dataset with 6 queried cell types highlighted in colour. (2) Spatial map of segmented single cells in ISS tissue section highlighting the queried cell type set. Inset shows raw microscopy images of DAPI and cell segmentation label masks that can be viewed on the app. (3) Visium tissue section with spot label masks. Inset shows raw H&E images and Visium spot label masks. (4) Cell set search console to cross-query multiple cell types across scRNA-seq and ISS datasets. (5) Cell type search console to cross-query a single cell type across all three datasets including Visium. (6) Gene search console to cross-query gene expression across all datasets. B. Cell type cross-query snapshot. Selecting chondroprogenitors via the cell type search console simultaneously highlights their cell cluster in scRNA-seq and their spatial locations in ISS and Visium datasets. On Visium data, the predicted abundance of chondroprogenitors per Visium spot is shown. C. Gene expression cross-query snapshot. Selecting the chondrocyte lineage marker COL2A1 via the gene search console returns its expression pattern in all three modalities, plotted per cells or Visium spots. D. Diagram of analysis workflow for the identification of spatially variable gene expression in chondroprogenitors. E. App snapshots showing enrichment of WNT5A expression in distal chondroprogenitors in ISS and Visium datasets. White dashed lines show the outline of developing bone where chondroprogenitors are located. ISS shows label masks of only chondroprogenitor cells. Insets show close up images of distal and proximal chondroprogenitors. F. App snapshot showing scRNA-seq and ISS datasets only. G. App snapshot showing scRNA-seq and Visium datasets only.

To showcase the capabilities of WebAtlas, we applied it to a multi-modal atlas of the developing human lower limb that integrates scRNA-seq, Visium and In Situ Sequencing (ISS) technologies (Fig 2A). The scRNA-seq and Visium datasets were generated previously^17^. To complement these with imaging-based ST data, we used a custom ISS pipeline to quantify the expression of 90 genes at single cell spatial resolution (see Methods). To facilitate the integration and comparison of different modalities, we curated ISS genes from the scRNA-seq atlas, focusing on cell type markers, and applied ISS to a tissue section adjacent to one of the Visium slices profiled in our previous study.

To harmonise lower limb atlas modalities, we computationally integrated the scRNA-seq reference with each ST dataset. The scRNA-seq and Visium integration, which deconvolves cell types in Visium data, was performed using Cell2location^9, 17^. To integrate scRNA-seq and ISS, we used StabMap^10^ to create a joint embedding, where we annotated cell types and imputed full transcriptomes of ISS-resolved cells based on the scRNA-seq reference data (Sup Methods, Sup Fig 1). Finally, we ingested all datasets via the WebAtlas pipeline to convert all elements, including raw images (*i.e.* Hematoxylin and Eosin (H&E) brightfield images for Visium and fluorescent microscopy for ISS) and cell/spot label masks from ST, into the Zarr format, and visualised them on the web portal.

WebAtlas enabled coordinated navigation of integrated transcriptomic datasets in the lower limb atlas (Fig 2A). We could readily cross-query cell types across all three modalities (Fig 2B). This facilitated comparison of spatial cell type locations between Visium and ISS datasets, where we observed consistent regional patterns for many cell types including chondrocyte and epithelial subtypes matching prior knowledge^17^ (Fig 2B, Sup Fig 2), validating our computational integration approach. Owing to its increased resolution, we could distinguish finer spatial cell type patterns in ISS than Visium, such as endothelial cells surrounding digits and a proximo-distal gradient of chondrocyte differentiation (Sup Fig 2). Beyond cell types, we could cross-query gene expression across the lower limb atlas (Fig 2C). Examining the scRNA-seq and ISS integration, we observed accurate imputed expression patterns of known marker genes that were missing from the ISS gene panel (Sup Fig 4).

We next used WebAtlas to investigate the spatial dynamics of limb development, focusing on the chondroprogenitor cells from which the cartilage anlage arises. Leveraging the imputed transcriptomes of ISS-resolved single cells, we sought to identify genes with spatially variable expression patterns within chondroprogenitors (Fig 2D). This identified novel proximo-distal expression gradients of chondrogenesis related genes. *WNT5A*, which induces chondrogenesis^18^, was upregulated in distal chondroprogenitors (Fig 2E), whereas cartilage extracellular matrix genes such as *MATN1, 3 & 4*^19^ were upregulated in proximal chondroprogenitors (Sup Fig 3). Importantly, we validated these expression gradients on adjacent Visium sections using WebAtlas (Fig 2E, Sup Fig 3). Our observations are consistent with the proximo-distal nature of limb maturation, with immature distal chondroprogenitors beginning chondrogenesis whilst more mature proximal cells actively secrete extracellular matrix proteins. Finally, WebAtlas also enabled us to flexibly visualise subsets of lower limb atlas modalities, such as scRNA-seq data paired with either ISS (Fig 2F) or Visium (Fig 2G).

To demonstrate WebAtlas compatibility with other spatial technologies, we applied it to Xenium, MERSCOPE and Visium CytAssist datasets as well as a mouse embryonic atlas integrating scRNA-seq and seqFISH^20^ (Sup Table 2, Sup Figure 4). WebAtlas identifies key data elements from these diverse inputs and generates a standardised output, demonstrating our technology-agnostic approach. WebAtlas is also scalable to large datasets, exemplified here on a scRNA-seq study with over 900,000 cells and spatial datasets with over 700,000 cells (MERSCOPE) and up to 1.1 million RNA molecules/spots (ISS) (Sup Table 2, Sup Figure 4). We provide interactive access to all 19 datasets used in this study, including the lower limb atlas, via the WebAtlas portal at https://cellatlas.io/webatlas.

Taken together, we provide an intuitive pipeline for interactive exploration of integrated single cell and spatial transcriptomic datasets. WebAtlas incorporates data from diverse atlassing technologies into easily accessible multi-modal tissue atlases, and leverages scalable, standardised and FAIR data formats. We demonstrate exploration of a multi-modal human lower limb atlas, and use WebAtlas to interpret new biological insights arising from computational integration of single cell and spatial data. A current limitation of WebAtlas, despite leveraging GPU-accelerated rendering, is the scaling of points (RNA molecules) (Sup Note 1). This can be addressed in the future by rasterization of point data or multi-scale point clouds.

WebAtlas will facilitate the creation of rich and easily accessible human tissue atlases for biologists. We envision diverse use cases in the community, including anatomical and pathology annotation of spatial datasets, data dissemination alongside publications, and centralised reference atlases all made possible by cooperation between community efforts such as NGFF and SpatialData, uniform data structures and reusable software. In the future, cloud-optimised reference datasets could be potentially hosted in central repositories, such as Bioimage Archive^21^ or Image Data Resource^22^ with the data served directly through external web apps or analysed on the fly^23, 24^, creating a globally accessible and flexible framework for multimodal tissue atlases.

## Data availability

All datasets presented on this paper, including the lower limb atlas, are publicly available for exploration through the WebAtlas portal at https://cellatlas.io/webatlas.

## Code availability

All software code has been made publicly available on Github at https://github.com/haniffalab/webatlas-pipeline. Each software release is permanently archived on Zenodo at https://doi.org/10.5281/zenodo.7405818. Comprehensive documentation, tutorials and sample workflows are available at https://haniffalab.github.io/webatlas-pipeline.

## Acknowledgements

We thank the authors of the human lower limb atlas study for sharing data, Sébastien Besson for consultation on use of OME-NGFF as a common format, Pavel Mazin for help with Cell2location analysis of the Lower limb Visium data and Sanger Institute’s Informatics and Digital Solutions team for infrastructure support for hosting the datasets and website, and Jason Swedlow for comments on the manuscript. This work was funded by Wellcome Trust core funding (reference: 206194 and 220540/Z/20/A) and Strategic Science Award (reference: 221052/Z/20/Z, 221052/A/20/Z, 221052/B/20/Z, 221052/C/20/Z, and 221052/E/20/Z) to M.H. and O.A.B., and a Wellcome Trust Senior Research Fellowship Award (223092/Z/21/Z) to M.H.. J.M. was supported for work on OME-NGFF by grant numbers 2019-207272 and 2022-310144 and on Zarr by grant numbers 2019-207338 and 2021-237467 from Chan Zuckerberg Initiative DAF, an advised fund of Silicon Valley Community Foundation, and was funded by the Deutsche Forschungsgemeinschaft (DFG, German Research Foundation) – 501864659 as part of NFDI4BIOIMAGE. S.G was supported for work on Stabmap imputation by Australian Research Council Discovery Early Career Researcher Award (DE220100964) funded by the Australian Government and Chan Zuckerberg Initiative Single Cell Biology Data Insights grant (2022-249319).

## Author Contributions

T.L., D.H., D.B.L., M.H. and O.A.B. conceived the study. T.L. developed the WebAtlas Pipeline and performed ISS image analysis. D.H. and D.B.L developed the WebAtlas Pipeline, App and processed the sample datasets. M.P. supported WebAtlas implementation. K.R. generated ISS data. E.T. collected limb tissue samples. J.E.G.L., P.H. and S.T. shared limb scRNA-seq and Visium data, and contributed to limb ISS data interpretation J.M. consulted on the OME-Zarr specifications and reviewed and edited the manuscript. S.G. performed ISS-scRNA-seq data integration and analysis. T.L., D.H., D.B.L., M.H. and O.A.B. wrote the manuscript with feedback from all authors.

## Supplementary Note 1: The landscape of platforms for dissemination and navigation of tissue atlas data

The integration of single cell and spatial transcriptomics has become a standard approach for building comprehensive tissue atlases. However, the diversity and scale of data types poses significant challenges to usable and interpretable access of multi-modal tissue atlases. Here, we provide a comparison of the WebAtlas pipeline to existing alternative platforms (Sup Table 1) and elaborate our key advances that democratise access to complex tissue atlases.

### Simultaneous browsing and cross-query of multiple modalities

The coordinated navigation of scRNA-seq and ST datasets greatly facilitates biological insights from tissue atlases. Amongst existing platforms, WebAtlas uniquely enables the browsing and cross-query of integrated scRNA-seq and ST modalities. While the MoBIE plugin supports browsing individual imaging-based ST datasets or overlaying images from correlative microscopy, WebAtlas allows simultaneous browsing of integrated single cell and spatial transcriptomic datasets as well as their cell type and gene expression cross-query. This is enabled by 1) the WebAtlas data ingestion pipeline that can load datasets from most common scRNA-seq and ST technologies, and configure them for integrated visualisation and 2) the Vitessce framework that supports visualisation of multimodal single cell and spatial datasets.

### Cloud-optimised data storage

WebAtlas adopts the cloud-ready Zarr format, which provides an agnostic approach to the underlying storage system, and employs an array chunking strategy. The data is stored as a collection of text files, consisting of the data divided into compressed chunks which can be read individually or in parallel, allowing very large datasets to scale more efficiently in the cloud. While TissUUmaps 3 and SODB use native AnnData objects for tabular data and TissUUmaps 3 uses OME.tif file format for imaging data, WebAtlas converts all scRNA-seq and ST data objects to Zarr. Our data standardisation is strongly aligned with the community-defined next-generation file format (NGFF)^15^ built on Zarr.

### Web application

WebAtlas does not require users to install specialised local software, unlike the Loupe Browser (10X Genomics) and the Fiji plugin MoBIE, and is easily accessible on a web browser. This is enabled through the adoption of the cloud-ready Zarr format and the Vitessce serverless web framework in the WebAtlas pipeline.

### Multiscale raster image visualisation

ST technologies generate complex tissue image data, ranging from brightfield H&E images in Visium to multi-cyclic and multi-channel fluorescent microscopy images in imaging-based ST methods, such as ISS. The diversity and scale (*i.e.* high resolution imaging of large tissue areas) of these image data types poses challenges to their online dissemination. While CellxGene and SODB platforms can serve tabular gene expression files, they do not support online browsing of full resolution tissue images. WebAtlas provides efficient multi-scale visualisation of diverse ST tissue image types via the OME-Zarr format and the Vitessce tool.

### Cell and RNA segmentation visualisation

The analysis of imaging-based ST data can segment single cells and RNA molecules/spots in tissues. While SODB is limited to visualising segmented single cells as points, WebAtlas uses OME-Zarr and Vitessce for multi-scale visualisation of cell segmentation label masks that can depict complex cell shapes and cell-cell contacts at high resolution *in situ*. WebAtlas also visualises RNA molecules as points embedded in AnnData.

### Limitations

We observed that while many millions of RNA molecules can be ingested into the WebAtlas Pipeline and that the WebAtlas App can visualise 1.1 million RNA molecules on the lower limb ISS dataset and 6.5 million molecules downsampled from a Xenium dataset (see examples on the WebAtlas portal), the responsiveness of the WebAtlas app is dramatically reduced when we render over 10 million molecules on Xenium and MERSCOPE datasets (not shown). Sitting underneath Vitessce is a visualisation framework called deck.gl (https://deck.gl/), that leverages the WebGL 2.0 standard (https://registry.khronos.org/webgl/specs/latest/2.0/) to access GPUs on the client device. This significantly improves the performance of visual exploration of large datasets in the web browser. However, despite the GPU-accelerated rendering in Vitessce, there is still eventually a limit to the number of points that can be rendered in the web browser, and the user experience will vary depending on the specification of the client device.

To overcome this scalability limitation, it is necessary to change the architecture of how point data is visualised. Possible future scalability improvements include the rasterization of point data so that point data can be rendered efficiently as multi-scale NGFF images. Whilst offering a visual representation of the data, this solution would be at the expense of the user being able to interact with the molecule data through the web interface. Alternatively, multi-scale point clouds would provide a sequence of progressively more downsampled copies of the point data. Using a similar concept to the multi-scale pyramidal image formats, the app would then only load the appropriate point resolution and region that corresponded to the area of interest in the sample that the user requested to view.

## Supplementary Methods

### Lower limb single cell RNA-seq and Visium data

We used 10X scRNA-seq and Visium data previously generated and analysed by Zhang et al. ^1^. The scRNA-seq dataset spans the embryonic limb from PCW5 to PCW9, and consists of 125,955 cells that were annotated to 60 cell types. The Visium data spans PCW5 to PCW8 across 8 capture areas/chips. The whole scRNA-seq and Visium datasets were integrated using Cell2location, and subsequently used in this study. In the integrated lower limb WebAtlas, we visualise the whole scRNA-seq dataset along with one Visium capture chip from a PCW5.5 donor that matches our ISS dataset.

The tabular gene expression and cell type annotation tables were originally formatted as AnnData, including the cell abundance estimates in the Visium dataset generated by Cell2location analysis. The H&E images of Visium data were formatted as raw tiff files and loaded from 10X SpaceRanger files.

### Lower Limb ISS data generation

#### Sample acquisition and ethics

The lower limb of a PCW5.5 human embryo was obtained from an elective termination with informed consent under REC 96/085 (East of England - Cambridge Central Research Ethics Committee). The limb was embedded in optimal cutting temperature medium (OCT) and frozen at −80°C on an isopentane-dry ice slurry. Cryosections were cut at a thickness of 10 μm using a Leica CM1950 cryostat and placed onto SuperFrost Plus slides (VWR).

#### Customised In situ sequencing pipeline

In situ sequencing was performed using the 10X Genomics CARTANA HS Library Preparation Kit (1110-02, following protocol D025) and In Situ Sequencing Kit (3110-02, following protocol D100), which comprise a commercialised version of HybISS^2^.

A limb section was fixed in 3.7% formaldehyde (Merck 252549) in PBS for 30 minutes, washed twice in PBS for 1 minute each, permeabilized in 0.1 M HCl (Fisher 10325710) for 5 minutes, and washed twice again in PBS, all at room temperature. Following dehydration in 70% and 100% ethanol for 2 minutes each, a 9 mm diameter (50 μl volume) SecureSeal hybridisation chamber (Grace Bio-Labs GBL621505-20EA) was adhered to the slide and used to hold subsequent reaction mixtures. Following rehydration in buffer WB3, probe hybridisation in buffer RM1 was conducted for 16 hours at 37°C. The 90-plex probe panel included 5 padlock probes per gene, the sequences of which are proprietary (10X Genomics CARTANA). The section was washed with PBS-T (PBS with 0.05% Tween-20) twice, then with buffer WB4 for 30 minutes at 37°C, and thrice again with PBS-T. Probe ligation in RM2 was conducted for 2 hours at 37°C and the section washed thrice with PBS-T, then rolling circle amplification in RM3 was conducted for 18 hours at 30°C. Following PBS-T washes, all rolling circle products (RCPs) were hybridised with LM (Cy5 labelling mix with DAPI) for 30 minutes at room temperature, the section was washed with PBS-T and dehydrated with 70% and 100% ethanol. The hybridisation chamber was removed and the slide mounted with SlowFade Gold Antifade Mountant (Thermo S36937). Imaging of Cy5-labelled RCPs at this stage acted as a QC step to confirm RCP (‘anchor’) generation and served to identify spots during decoding. Imaging was conducted using a Perkin Elmer Opera Phenix Plus High-Content Screening System in confocal mode with 1 μm z-step size, using a 63X (NA 1.15, 0.097 μm/pixel) water-immersion objective and 7% overlap between adjacent tiles. Channels: DAPI (excitation 375 nm, emission 435-480 nm), Atto 425 (ex. 425 nm, em. 463-501 nm), Alexa Fluor 488 (ex. 488 nm, em. 500-550 nm), Cy3 (ex. 561 nm, em. 570-630 nm), Cy5 (ex. 640 nm, em. 650-760 nm).

Following imaging, the slide was de-coverslipped vertically in PBS (gently, with minimal agitation such that the coverslip ‘fell’ off to prevent damage to the tissue). The section was dehydrated with 70% and 100% ethanol, and a new hybridisation chamber secured to the slide. The previous cycle was stripped using 100% formamide (Thermo AM9342), which was applied fresh each minute for 5 minutes, then washed with PBS-T. Barcode labelling was conducted using two rounds of hybridisation, first an adapter probe pool (AP mixes AP1-AP6, in subsequent cycles), then a sequencing pool (SP mix with DAPI, customised with Atto 425 in place of Alexa Fluor 750), each for 1 hour at 37°C with PBS-T washes in between and after. The section was dehydrated, the chamber removed, and the slide mounted and imaged as previously. This was repeated another five times to generate the full dataset of 7 cycles (anchor and 6 barcode bits).

### Lower limb ISS image data processing

#### 1. Image pre-processing

We used proprietary software provided by Perkin Elmer for the initial processing of raw ISS image data. This entailed illumination correction, maximum Z intensity projection and stitching, resulting in the generation of an ome.tiff file per imaging cycle that encompasses all the channels (DAPI, Atto425, Alexa Fluor 488, Cy3 and Cy5).

#### 2. Image registration

We used the Microaligner package for a two-step registration process of ISS imaging cycles^3, 4^. The first step is Affine feature-based registration, where the DAPI channel in the first ISS cycle serves as the reference image and the subsequent cycles are registered to this reference. We begin by detecting image features in the DAPI channels using the FAST feature point finder algorithm in OpenCV package^5^, which identifies image areas with significant intensity changes. Next, the DAISY feature descriptor algorithm extracts histograms of oriented gradients for each identified feature point. The extracted feature points are then matched using the FLANN-based KNN matcher algorithm across cycles, which determines the correspondence between the features of the reference and moving images. The matches are filtered based on the default distance threshold between neighbouring features, and the resulting matched feature coordinates are aligned using the RANSAC algorithm in OpenCV to compute the affine transformation. The process is applied to tiled images with tile size of 6000 by 6000 pixels to optimise the alignment and reduce memory usage. For each tile, a transformation matrix is derived after applying the DoG function with predefined kernel sizes, which is eventually unified by employing the *matmul* function in Python.

The second step is non-linear optical flow-based registration that relies on the Farneback method (calcOpticalFlowFarneback) in OpenCV that identifies pixels with the highest similarity within a given window, with fine tuned parameters - pyr_scale=0.5, levels=3, winsize=51, iterations=3, poly_n=1, poly_sigma=1.7, flags=OPTFLOW_FARNEBACK_GAUSSIAN. For each pixel, the method computes a 2D vector that characterises the movement of the pixel from one image to the other. This is applied to tiled images with a tile size of 1000 by 1000 pixels with an overlap of 100 pixels between adjacent tiles.

#### 3. ISS barcode decoding with PoSTcode

To decode individual RNA transcripts from cyclic ISS images, we used the PoSTcode barcode decoding algorithm^3^ and customised image preprocessing. To improve the accuracy of downstream spot-calling and quality of intensity-based decoding, we first applied the white hat filter with the kernel size of 5 pixels to filter out noise from all coding channels. Subsequently, the transcript detection was performed exclusively on the anchor channel using the ‘locate’ method in TrackPy^6^ with percentile equal 90, spot size equals 5 and separation equals 4. One intensity profile for each transcript was extracted from the registered image generated in the last step. This intensity profile is of shape 4×6 representing the intensity extracted from 4 channels per cycle and from all the 6 cycles. Yet, to improve decoding outcome, we expanded the searching range of the maximum intensity to +/− 2 pixels across coding channels. The decoding step in PoSTcode takes this 4×6 matrix and the codebook from CARTANA as input and returns prediction of gene type for each transcript with a confidence value. Only transcripts with a value higher than 0.97 were kept and saved as a .csv file for downstream processing.

### 4. Single cell segmentation

To segment single cells from the registered image stack, we applied the cell segmentation in CellPose^7^ using the pretrained ‘cyto2’ model on DAPI channel with the cell size of 70 pixels in diameter. To mimic the cytoplasm boundary, expansion of 10 pixels is applied and the expanded cell segmentation was used to generate the cell by gene expression matrix. Due to the large memory requirements, we adopted a strategy of dividing the whole images into smaller tiles and performed the segmentation on each of the tiles individually. Following this, we stitched the tiles back together to reconstruct the complete image without compromising much segmentation accuracy. There were in total 117,788 cells detected.

#### 5. Anndata object generation

The decoded 1,164,802 spots were assigned to the 117,788 cells using the STRtree^8^. Out of the 117,788 cells, only 66,675 cells were kept after filtering out cells with less than 4 transcripts. The output is saved as an AnnData object.

### Lower limb ISS and scRNA-seq data integration and analysis

#### 1. Generating reference embeddings for scRNA-seq and ISS

For the scRNA-seq data, we used the principal components from the previous study as the target reference embedding for mosaic integration. For the ISS data, we selected cells with at least 5 detected transcripts in at least 3 genes. We then calculated additional spatially-informed features beyond per-cell expression counts by calculating the gene counts from among the nearest 25 transcripts detected from the centroid of each ISS-resolved cell. These new features were concatenated among the per-cell expression counts and converted to log-counts. We calculated 30 principal components and treated this as the reference embedding for ISS.

#### 2. Mosaic integration using StabMap

Using the scRNA-seq and ISS reference embeddings, we jointly mapped these data onto each other using StabMap, which included a rescaling of embedding values according to L1 norm. Then to ensure no residual modality-specific effect, we performed horizontal data integration using Harmony. This resulted in a joint corrected StabMap lower-dimensional embedding for all cells.

#### 3. Imputation and Cell type classification

We used the joint embedding for all cells to perform point imputation and cell type classification. For point imputation, for each ISS-resolved cell, we calculated the mean logcount gene expression across the five nearest euclidean distance scRNA-seq resolved cells within the corrected lower-dimensional embedding. For cell type classification, we used the K-nearest neighbours algorithm, with K = 5, and selected the majority class for each ISS-resolved cell, with ties broken by the classes nearest to the ISS-resolved cell. To denoise individual cell classifications, we reassigned cell type labels of each ISS-resolved cell to the majority class of the ISS-resolved cell’s 30 most proximate cells in euclidean distance.

#### 4. Spatially variable gene expression analysis

We next performed spatially variable gene expression analysis on the imputed and cell type classified ISS data. We downsampled to 1,000 cells within ChondroProg, randomly sampled across the entire ISS spatial coordinates. We then used scHOT^9^ to measure departure from homogenous expression across space via weighted means for each cell, with weights proportional to euclidean distances in space spanning 5% of the nearest cells. The scHOT observed test statistics were used to rank imputed genes according to departure from homogenous spatial expression, and the top 20 imputed genes were identified as spatially differentially expressed.

### WebAtlas data ingestion

The WebAtlas data ingestion pipeline requires the user to provide a YAML file that defines input datasets. Each dataset can be composed of tabular data and/or images. Currently supported dataset types are AnnData object (*e.g.* HDF5 files of tabular gene expression from scRNA-seq and ST), Visium SpaceRanger output (up to version 1.2.0), Xenium output (up to version 1.3), MERSCOPE output (version 2022.5.26), molecules CSV/TSV file (*e.g.* RNA spots from customised ISS), and any raster images including raw microscopy and cell/spot label masks supported by bioformats2raw. The ingestion of different scRNA-seq and ST modalities is detailed below.

The user must specify the paths or URLs for each dataset and their corresponding types, along with visualisation options for Vitessce, including the final URL hosting the data. We provide various template YAML files for different modalities on our Github repository (see Code availability section).

The pipeline converts tabular AnnData to Zarr format via the canonical write_zarr function within the ScanPy package^10^. Raster images are converted to OME-Zarr using the bioformats2raw tool using default parameters.

The pipeline outputs data objects converted to Zarr and a View Config JSON file, which configures the WebAtlas web application (see below). To visualise datasets on the web app, the user needs to ensure that the Zarr data objects and the View Config file are accessible, which can be accomplished through local hosting or by placing the files onto cloud-based services such as AWS S3 bucket or Google cloud (see Vitessce guidance at http://vitessce.io/docs/data-hosting/).

#### 1. scRNA-seq data

We ingest scRNA-seq datasets in the AnnData format and convert them to AnnData-Zarr.

#### 2. Visium data

We can ingest raw Visium and Visium CytAssist datasets from the SpaceRanger output directories. We convert tabular spot by gene expression files to AnnData-Zarr and H&E raster images to OME-Zarr. To visualise Visium spots to be overlaid on the H&E images, we generate label images of Visium spots based on the spatial information included in the SpaceRanger output files defining each spot’s centre coordinates and diameter. For Visium datasets that have been integrated with scRNA-seq by Cell2location, we can ingest Cell2location output AnnData objects that list the deconvolved cell type abundances per Visium spot. To visualise deconvolved cell types on Visium data, which are formatted as continuous cell abundance numbers per Visium spot, further preprocessing is required and is described in the integrated modality visualisation section below.

#### 3. Custom ISS data

Our ISS image analysis pipeline generates 1) tabular cell by gene expression files that are loaded as AnnData-Zarr and 2) raster images of raw microscopy data and segmentation label masks that are loaded as OME-Zarr. Additionally, if available, segmented RNA spots/molecules can be loaded as embedded in the AnnData format.

#### 4. Xenium data

We used 4 Xenium datasets provided by 10X Genomics, including a human breast tumour^11^ (Xenium file format version 1.0.1) and human brain tissue including glioblastoma tumours (Xenium file format version 1.3.0) (provided in https://www.10xgenomics.com/resources/datasets/xenium-human-brain-preview-data-1-standard). The tabular cell by gene expression input files, available as 10x-Genomics-formatted HDF5 files, are ingested and converted to AnnData-Zarr using a dedicated loading function in ScanPy. Raster microscopy images are ingested to OME-Zarr. The cell segmentation masks are formatted as polygons in the Xenium file format, and are converted to label images in OME-Zarr format. Additional information provided by Xenium technology, such as cell centroid coordinates and default clustering labels, are loaded using additional scripts in the pipeline.

#### 5. MERSCOPE data

We used a human breast cancer MERSCOPE FFPE dataset released by Vizgen (Human Immuno-oncology Data Release from https://vizgen.com/data-release-program/). The MERSCOPE data format stores metadata in separate files, necessitating the development of specialised loading functions to construct the AnnData object as well as to generate labelled images. We use the pandas package to load the multiple CSV files from the MERSCOPE output. We load the cell by gene matrix filtering out blank control barcodes which we identify by being prefixed with “Blank”. We load cell metadata and along with the expression matrix we build an AnnData object. To be able to map cells to labelled images we transform the cell centroid micron coordinates included in each cell metadata with a transformation matrix provided by MERSCOPE to obtain pixel coordinates. In a similar manner, we obtain segmentation pixel coordinates from a cell boundaries HDF5 file and the micron to pixel transformation matrix. We use these segmentations to generate labelled images in tiff format, where each segmentation is assigned the corresponding cell ID. As for the raw image, we aim to obtain a single multipage tiff image that will then get converted to OME Zarr. MERSCOPE outputs each channel as a different tiff file and thus we first concatenate them into a single file through the pyvips^12^ package and set all necessary OME metadata that is then used by bioformats2raw when performing the conversion to Zarr.

### WebAtlas visualisation via Vitessce

The WebAtlas Data ingestion pipeline creates a Vitessce View Config JSON file to facilitate data visualisation, outlining pertinent information such as input datasets, specifications of each data type, embeddings to be represented, component layout within the app, and the transformed dataset’s behaviour in Vitessce. The View Config file conforms to the guidelines outlined in the Vitessce documentation (http://vitessce.io/docs/view-config-json/).

### WebAtlas visualisation of integrated modalities

To facilitate the integrated visualisation of gene expression and cell types in the scRNA-seq/ISS/Visium datasets, it is necessary to preprocess all the data.

Firstly, we manipulate the expression matrices of all data modalities to facilitate the visualisation of deconvolved cell type abundances in Visium. In the Visium data, we concatenate the cell type abundance predictions from Cell2location into the spot by gene expression matrix and identify which features are cell type predictions using a boolean column labelled “is_celltype”. The genes from the original spot by gene part of the matrix are then labelled as “is_gene.” This manipulation allows for the display of continuous cell type predictions generated by Cell2location, rather than showing only a single cell type prediction per Visium spot. Along with this, we expand the expression matrices of the corresponding scRNA-seq and ISS data to accommodate the “is_celltype” and “is_gene” columns and enable simultaneous searching across all modalities through the “featureList” component in Vitessce.

This expansion involves translating the original categorical cell type values in the scRNA-seq and ISS label encoding into a one-hot encoding matrix. This is done by representing each label as a binary vector with a value of 1 in the corresponding category and 0s elsewhere. The axes between all the modalities are intersected, and the objects are sliced to contain only these values. We also ensure that each observation (cell/spot) of each modality has a unique identifier - as an integer - across all modalities by adding offsets to different datasets cell/spot IDs. This is done because an overlap can cause incorrect visualisation in Vitessce. The intersected AnnData objects then get written into AnnData-Zarr.

Secondly, in the View Config file, an appropriate coordination value is assigned for the type of observation for all three modalities, which may be cells or spots depending on the sequencing method used. The feature type and their corresponding values, which might be gene expression or cell type abundance, is also defined. These coordination values are used by Vitessce when the web application is rendered, and are leveraged to achieve the integrated visualisation. Users can utilise the cell type or gene lists to search across the chosen ontology and visualise the expression of selected features across all modalities, allowing visualisation of gene expression or cell type abundance across both spatial profiles and embedding spaces such as UMAP or t-SNE. This is also true when visualising hierarchical observations such as cell type or any other shared ontology. Additionally, we coordinate other properties such as the colour map range and zoom values, that can be controlled separately by dataset or in a unified manner, to emphasise or compare between modalities.

We manually create a Vitessce View Config file to enable the loading of gene and cell type subsets of the concatenated matrices through specific coordination values. Within the View Config file we include each modality as a separate dataset object. For each modality we define three observation-by-feature matrices. The first observation-by-feature matrix points to the concatenated genes and cell types matrix within the AnnData-Zarr. We set the “featureType” coordination value of this matrix as “combined”. This first matrix must be included so the software can access the full matrix. The second and third observation-by-feature matrices point to the same concatenated matrix but are filtered through the “featureFilterPath” option. We filter the matrices by pointing this “featureFilterPath” to the column within the AnnData object’s feature axis that contains the boolean values that indicates whether that feature is a gene or a cell type. We respectively specify the “featureType” coordination value to “gene” in one observation-by-feature matrix and “celltype” in the other. These filtered matrices allow us to then set controls that load only one subset of the concatenated matrix at a time. For the datasets’ image data we set the “featureType” coordination value as “combined”. We then use the three “featureType” values, “combined”, “gene” and “celltype”, in the coordination space of the View Config. We refer to the “combined” value within the layout definition for the scatterplot and spatial components. Distinctly, we define two feature list components and refer one to the “gene” “featureType” and the other to “celltype”. Thus, each list displays only the values that correspond to each feature type, and selecting a feature loads the respective column from the concatenated matrices which are visualised on the scatterplot and spatial components. The feature list components can load the complete set of features from any of the datasets as they contain the same data from the intersection step. For future visualisations that are similar, such View Config files can be used as a template to generate relevant configurations and examples of these are provided in the WebAtlas Github repo.

---

**Supplementary Figure 1.**
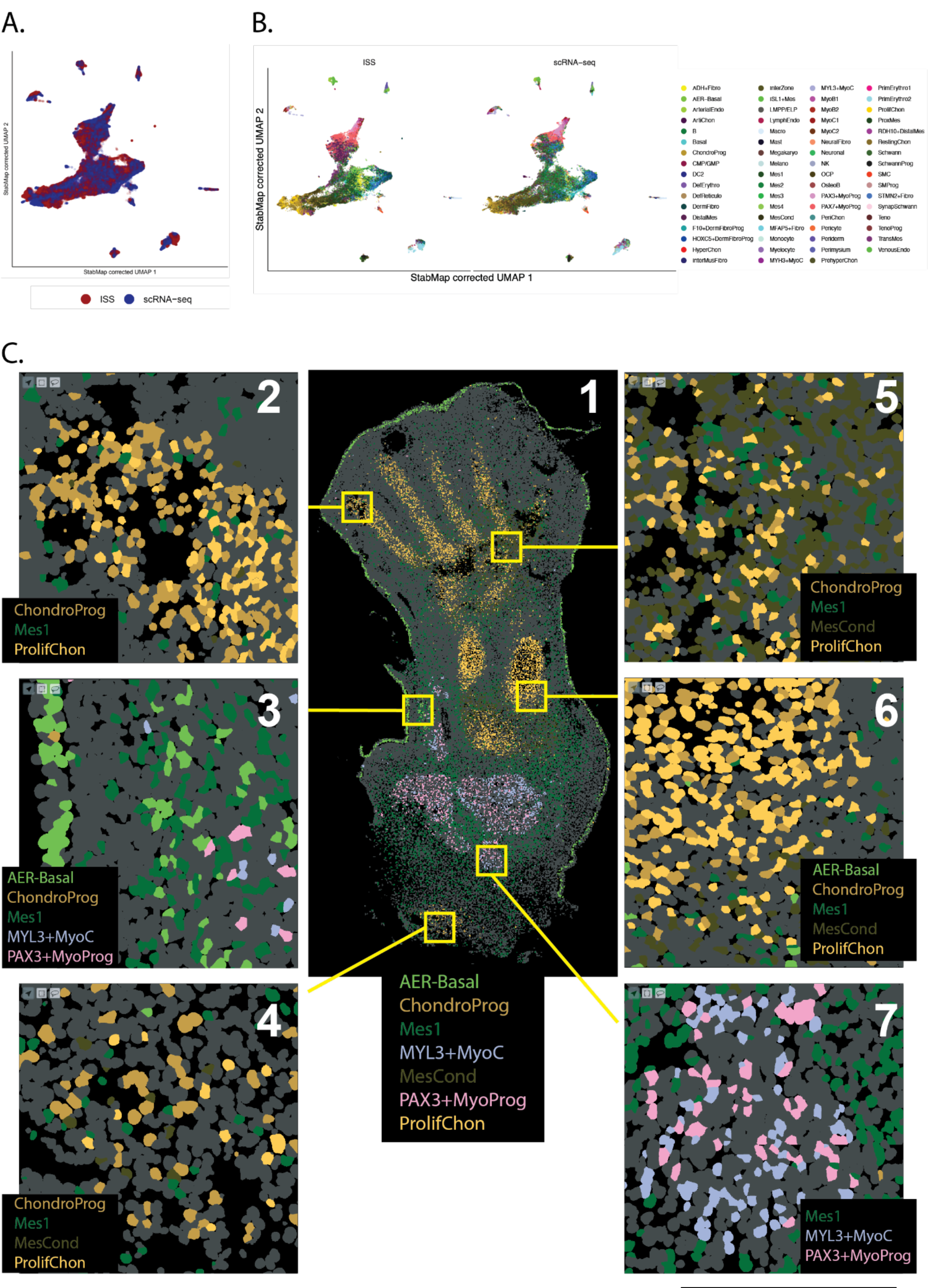
Integration of human lower limb scRNA-seq and ISS datasets. A) StabMap corrected UMAP displaying cells from scRNA-seq reference and ISS dataset. B) StabMap corrected UMAP displaying reference cell type annotations on the scRNA-seq dataset and the transferred cell type annotations on ISS dataset. C) Spatial map of annotated cell types in the lower limb ISS dataset shows expected regional patterns, validating StabMap computational integration. Panel 1; overview of the whole limb section displaying 7 out of 60 cell types. Panel 2; developing fifth digit consisting of chondroprogenitors (ChondroProg) and proliferating chondrocytes (ProlifChon). Inset 3; posterior aspect of the developing zeugopod, showing basal cells (AER-Basal) of the developing skin and mesenchymal progenitors (Mes1). Inset 4; limb-trunk junction with chondroprogenitors and proliferating chondrocytes. Inset 5; developing tarsal region showing mesenchymal condensate (MesCond), chondroprogenitors and proliferating chondrocytes. Inset 6; developing tibia consisting of mesenchymal condensate, chondroprogenitors and proliferating chondrocytes. Inset 7; Developing muscle of the stylopod, consisting of PAX3-expressing myoprogenitors (PAX3+MyoProg) and MYL3-expressing myocytes (MYL3+MyoC).

**Supplementary Figure 2.**
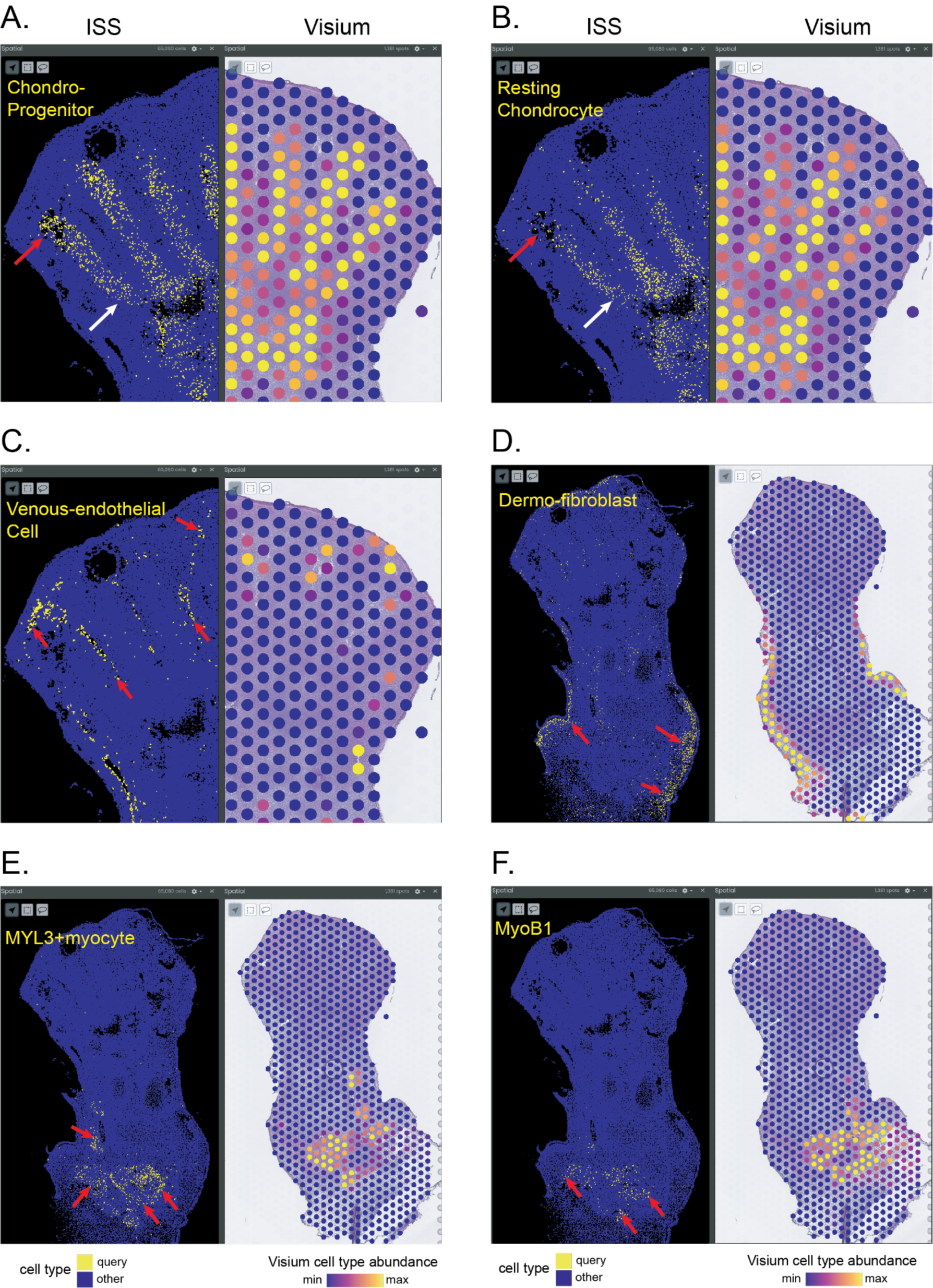
Spatial locations of cell types in ISS and Visium datasets. Panels show WebAtlas app snapshots of select cell types across different limb regions. ISS panels show the spatial maps of cell types annotated by StabMap computational integration and Visium panels show spatial cell type maps deconvolved by Cell2location computational integration. A,B) Chondroprogenitors and the more differentiated resting chondrocytes are mapped to the developing digits on both modalities. Chondroprogenitors are more abundant in the distal (indicated with red arrow) than the proximal parts (white arrow) of the digits, whereas resting chondrocytes show the inverse pattern, as more clearly distinguished by ISS data. C) The fine spatial localisation of venous-endothelial cells to interdigital areas (red arrows) is more clearly resolved in ISS than Visium data. D) Dermal fibroblasts are mapped to the developing skin in both modalities. E, F) MYL3+ myocytes and embryonic myoblasts (MyoB1) are mapped to the developing muscle in both modalities.

**Supplementary Figure 3.**
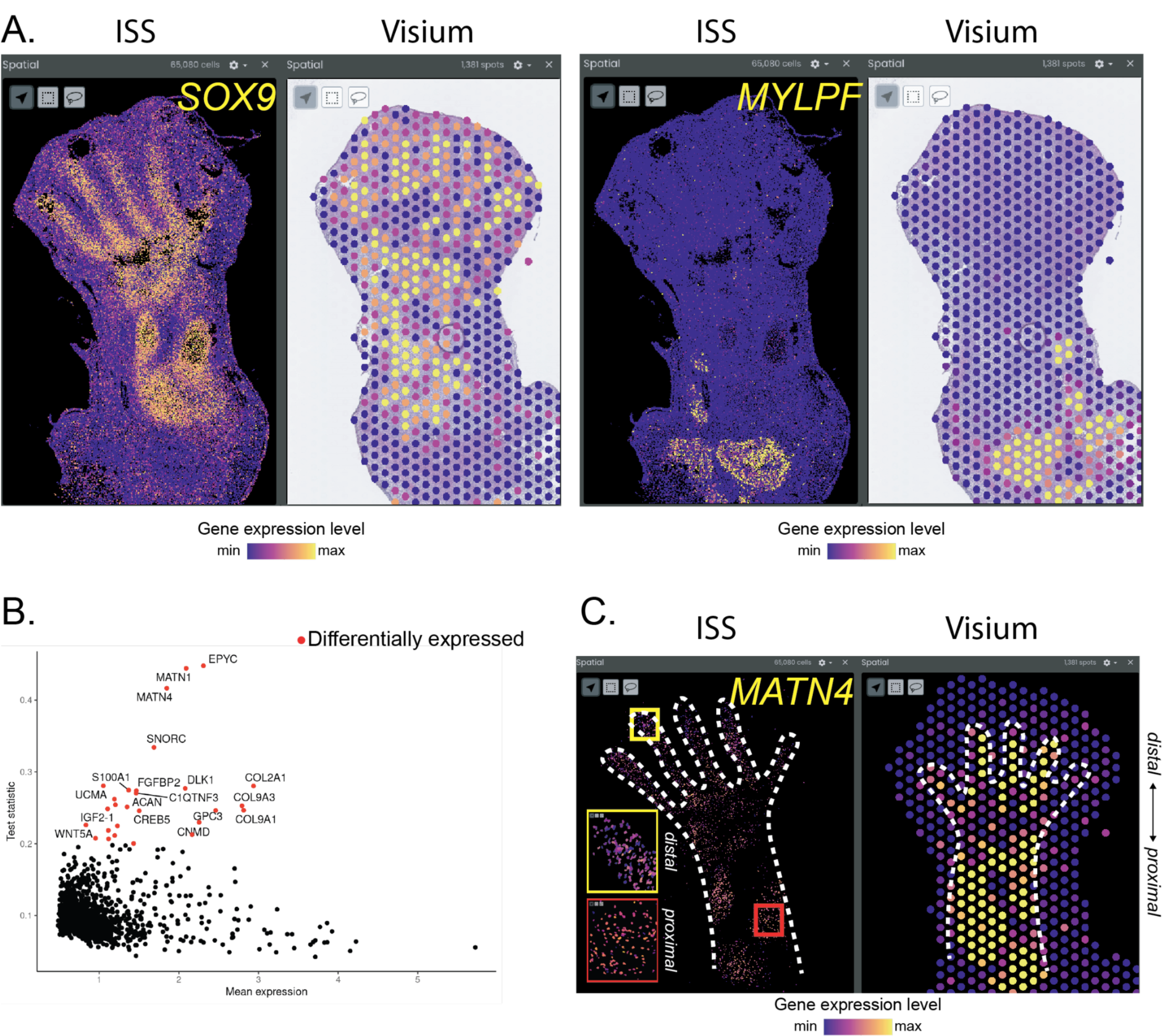
Imputation of unobserved gene expression and spatially variable gene expression analysis in the human lower limb. A) WebAtlas app snapshots showing spatial expression of SOX9 (chondrocyte lineage marker) and MYLPF (muscle marker) imputed in ISS versus observed in Visium. The imputed gene expression patterns of both markers are consistent with the known spatial patterns of their respective cell types and the observed expression in Visium. B) Scatter plot of mean imputed expression and scHOT observed test statistic for spatial differential expression for each imputed gene in the ChondroProg cell type in human lower limb. Top ranked genes are labelled. C) App snapshots showing enrichment of MATN4 expression in proximal chondroprogenitors in ISS and Visium datasets. White dashed lines show the outline of developing bone where chondroprogenitors are located. ISS shows label masks of only chondroprogenitor cells. Insets show close up images of distal and proximal chondroprogenitors.

**Supplementary Figure 4.**
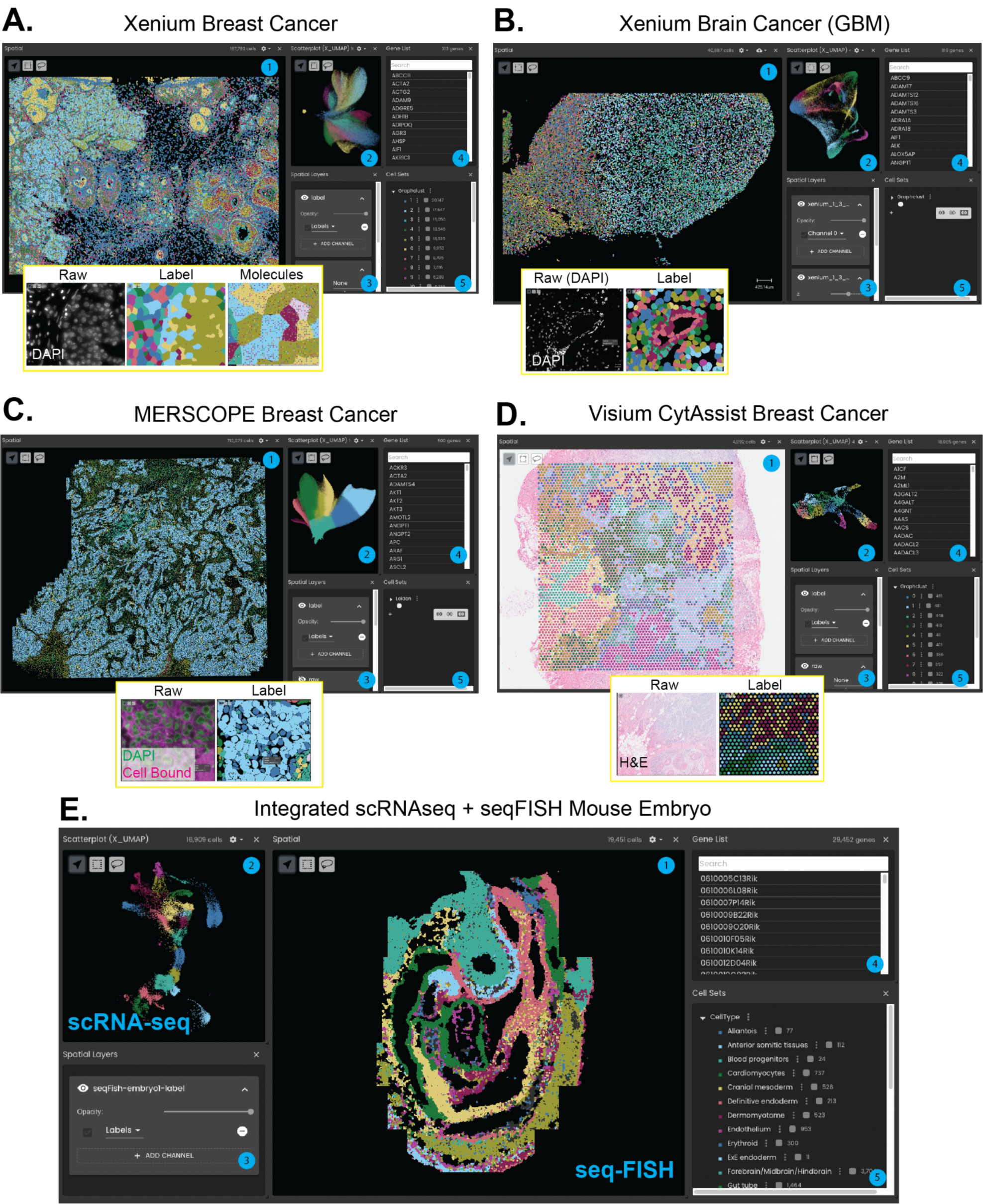
Different ST technologies on WebAtlas. WebAtlas app snapshots of (A) Xenium Human Breast Cancer, (B) Xenium Human Brain Cancer (Glioblastoma), (C) MERSCOPE FFPE Breast Cancer, (D) Visium CytAssist Human Breast Cancer, and (E) integrated scRNA-seq and seqFISH mouse embryo datasets. For each panel, the Vitessce component windows show the following as numbered. (1) Spatial map of segmented single cells or Visium spots coloured according to annotated cell types or cell/spot transcriptomic clusters. (2) UMAP representation of cell types or cell/spot transcriptomes. (3) Spatial layer console to toggle and adjust raster images, label masks and molecules. (4) Gene search console. (5) Cell type or cell/spot cluster search console. Inset panels show raw fluorescent or brightfield microscopy images, including DAPI for Xenium, DAPI and cell boundary staining for MERSCOPE, and H&E images for Visium CytAssist, as well as cell segmentation label images. The Xenium inset also shows RNA molecules.

**Supplementary Table 1:**
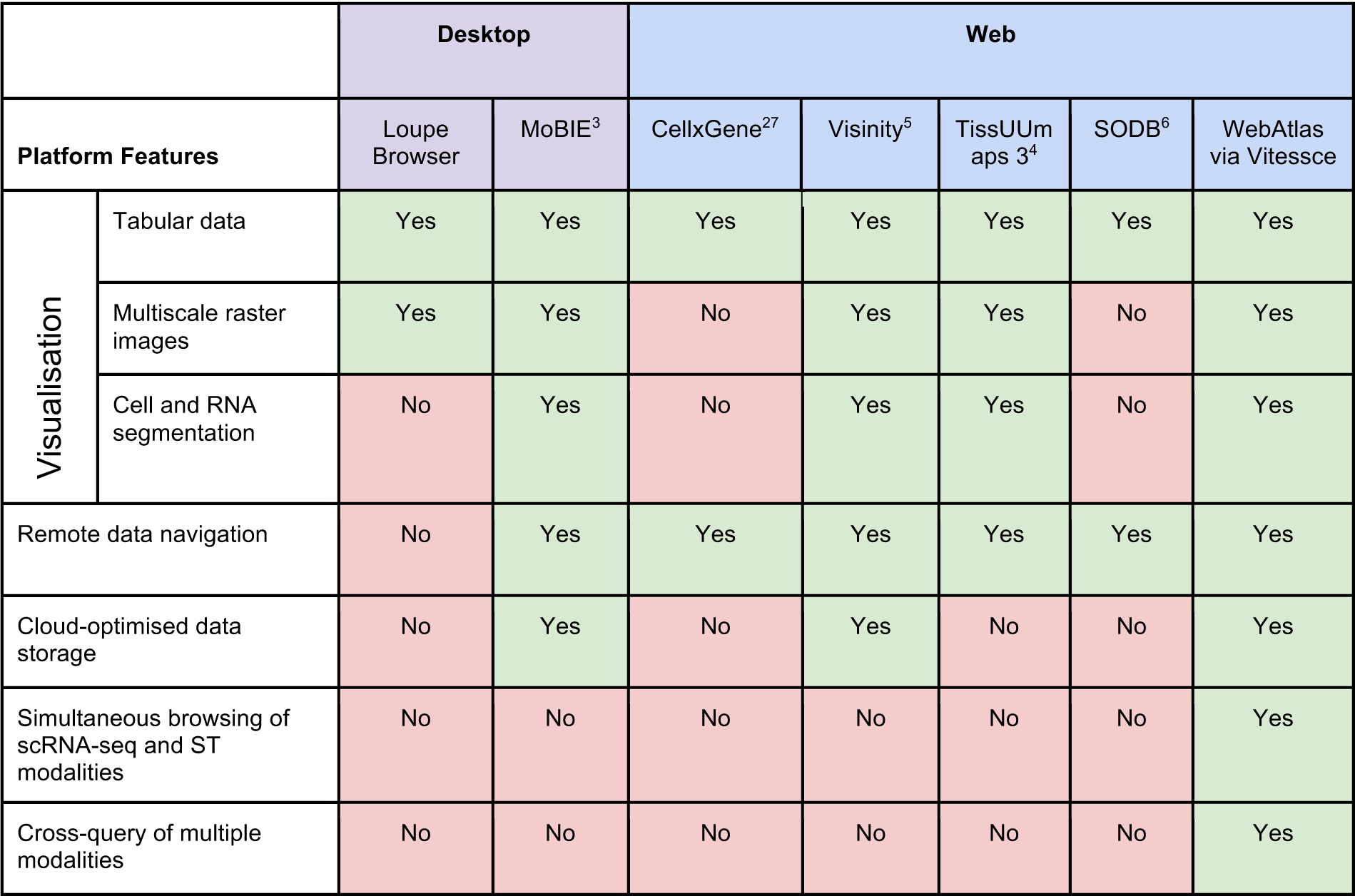
Comparison of WebAtlas with alternative platforms.

**Supplementary Table 2.** Datasets visualised on WebAtlas in this study and their specifications.

| Dataset and Reference | Technology | Cells/Visium spots | Image Size (pixels) | Genes | Molecules |
| --- | --- | --- | --- | --- | --- |
| Human Lower Limb<sup>17</sup> | scRNA-seq | 125,955 | N/A | 26,509 | N/A |
|  |  |  |  | 12,903 after multimodal intersection |  |
| Human Lower Limb (this study) | ISS | 65,080 | 26,525 x 47,831 | 90 | 1,164,802 (loaded on WebAtlas for the individual ISS dataset) |
|  |  |  |  | 26,522 after imputation |  |
|  |  |  |  | 12,903 after multimodal intersection |  |
| Human Lower Limb<sup>17</sup> | Visium* | 1,279 | 15,040 x 26,680 | 13,730 | N/A |
|  |  |  |  | 12,903 after multimodal intersection |  |
| Human Breast Cancer | Visium CytAssist | 4,992 | 19,505 x 21,571 | 18,085 | N/A |
| Human Breast Cancer<sup>25</sup> | Xenium | 167,782 | 35,416 x 25,779 | 313 | 43,664,540 (not loaded) |
| Human Brain Tumour | Xenium** | 40,887 | 39,794 x 23,900 | 319 | 13,324,742 (not loaded) |
| Human Breast Cancer | MERSCOPE | 710,073 | 110,485 x 94,805 | 500 | 490,398,542 (not loaded) |
| Foetal Immune<sup>26</sup> | scRNA-seq | 911,873 | N/A | 33,538 | N/A |
| SeqFISH Mouse embryo<sup>20</sup> | SeqFISH | 19,451 | N/A | 351 | 4,396,076 (not loaded) |
|  |  |  |  | 29,452 after imputation |  |
| SeqFISH Mouse embryo<sup>20</sup> | scRNA-seq | 16909 | N/A | 29,452 | N/A |

*All datasets can be publicly accessed on the WebAtlas portal.*
*\*The lower limb study profiled multiple donors and anatomical regions across 8 Visium chips (i.e. capture areas). One Visium chip with a PCW5.5 section was used for the integrated lower limb WebAtlas. The rest of the samples can be accessed as individual datasets in our portal.*
*\*\*Two additional human brain Xenium datasets can be accessed in our portal including healthy brain and Alzheimer's disease tissue sections.*

